# Intermuscular coherences of plantarflexors during walking suggest distinct neural origin and function for alpha and beta/low-gamma bands after stroke

**DOI:** 10.1101/2023.07.21.550018

**Authors:** CC Charalambous, MG Bowden, JN Liang, SA Kautz, A Hadjipapas

**Author notes:** Corresponding Author: *Charalambos C. Charalambous, PhD* Department of Basic and Clinical Sciences & Center for Neuroscience and Integrative Brain Research University of Nicosia Medical School, Nicosia, Cyprus 21 Ilia Papakyriakou, Block C, Rm 202 CY-1700, Nicosia, Cyprus ORCID: 0000-0002-1547-1839.

## Abstract

Plantarflexors provide propulsion during walking (late stance) and receive input from both corticospinal tract (CST) and corticoreticulospinal tract (CReST). Both descending motor tracts exhibit some frequency-specificity, which allows potential differentiation of neural drive from each tract using intermuscular coherence (IMC). Stroke may differentially affect each tract, thus impair the function of plantarflexors. However, the evidence concerning this frequency-specificity and its relation to plantarflexors’ neuromechanics post-stroke remains very limited. Here, we investigated the intermuscular coherences of alpha, beta, and low-gamma bands between the Soleus (SOL), Lateral Gastrocnemius (LG), and Medial Gastrocnemius (MG) muscles and their relationships with walking-specific measures (propulsive impulse; speed). Fourteen individuals with chronic stroke walked on a treadmill at self-selected and fast walking speed (SSWS and FWS, respectively). Inter-limb IMC comparisons revealed that beta LG-MG (SSWS) and low-gamma SOL-LG (FWS) IMCs were degraded on the paretic side. At the same time, within each limb, the IMCs, which were significantly different to a surrogate dataset denoting random coherence, were in the alpha band (both speeds). Further, alpha LG-MG IMC was positively correlated with propulsive impulse in the paretic limb (SSWS). Findings suggest differential functional role of alpha and beta/low-gamma, which may be related to the frequency-specificity of the underlying descending drives. The persistence of alpha in plantarflexors and its strong positive relationship with propulsive impulse suggests relative preservation and/or upregulation of CReST. Future research should address whether entraining motor system at alpha frequencies via neuromodulation can improve the neuromechanical function of paretic plantarflexors and subsequently promote post-stroke walking recovery.

**Key Points Summary:**

- Cortical and subcortical motor drives may be frequency-specific, have a role in walking, and be degraded after stroke.
- Whether this frequency-specificity exists and how it is related to neuromechanical function of ankle plantarflexors post-stroke remains to be determined.
- Here, we investigated bilaterally the intermuscular coherences of alpha, beta, and low-gamma bands for the Soleus (SOL), Lateral Gastrocnemius (LG), and Medial Gastrocnemius (MG) muscles and their relationships with walking-specific measures (propulsive impulse; self-selected and fast speed) during treadmill walking in individuals post-stroke.
- The beta LG-MG (self-selected speed) and low-gamma SOL-LG (fast speed) were degraded on the paretic side.
- Alpha coherence was significantly present across plantarflexors mainly on the non-paretic side (both speeds).
- Paretic alpha LG-MG was positively correlated with paretic propulsive impulse (self-selected speed).
- Given that paretic propulsive impulse is impaired post-stroke, entraining the motor system at alpha frequency via neuromodulation may improve propulsive impulse and subsequently promote post-stroke walking recovery.

## INTRODUCTION

Both cortical and subcortical motor drives contribute to walking (Drew, Prentice, and Schepens 2004; Drew and Marigold 2015; Nielsen 2002; Barthelemy et al. 2011). The cortical motor drive is derived from the corticospinal tract (CST); a pathway that is monosynaptic (i.e., corticomotoneuronal), fast, and responsible for the activation of the contralateral musculature (mainly distal muscles) during individuated and dexterous actions (Nathan, Smith, and Deacon 1990; York 1987; Lemon 2008). The subcortical motor drive is mainly derived from the corticoreticulospinal tract (CReST); a pathway that is oligosynaptic (multiple synapses between cortex and motoneuronal pool), slow, and responsible for the bilateral activation of muscles (mainly axial and proximal muscles) during gross movements (Peterson 1979b; Peterson 1979a; Peterson, Pitts, and Fukushima 1979; Jang and Lee 2019). During walking, CST is mainly responsible for the execution of proactive (i.e., adaptive) and reactive gait modifications (Drew and Marigold 2015; Barthelemy et al. 2011), whereas CReST is mainly responsible for the modulation of the coordination between limbs within each gait cycle and for the production of coordinated postural responses (Matsuyama et al. 2004).

After stroke, walking is one of the motor activities that is limited (Langhorne, Bernhardt, and Kwakkel 2011) with most individuals with stroke experience walking deficits, which may be lasting and with minimal recovery (Jorgensen et al. 1999). Depending on lesion characteristics (e.g., size, location), either pathway can be degraded after stroke (Paul et al. 2023) and can contribute to these neuromechanical and behavioral walking deficits (Srivastava et al. 2022). Therefore, better characterization of the consequences of functional and anatomical disruptions of both pathways after stroke can likely elucidate underlying mechanisms, which can be used as predictors of and targets for locomotor recovery after stroke.

Often the descending, motor drive can be characterized using corticomuscular coherence, or more commonly, intramuscular coherence (within the same muscle) and intermuscular coherence (IMC; between muscles) (Mima and Hallett 1999; Grosse, Cassidy, and Brown 2002). IMC quantified in this way captures rhythmic aspects of descending drive, which is common (coherent) between the pair of muscles investigated, and may be partly related to the rhythmic activity of the brain regions themselves where these pathways originate (Buzsáki and Draguhn 2004; Schnitzler and Gross 2005). Interestingly, there is some evidence to suggest that this descending drive indeed exhibits some frequency-specificity (Charalambous and Hadjipapas 2022). There are anatomical and physiological considerations, which support some frequency-specificity in the descending pathways. CST is a fast-tract involving a single synapse (Lemon 2008; Nathan, Smith, and Deacon 1990; York 1987), thus the time delay of this cortical drive is expected to be short (something which is indeed evident in short latencies of the contralateral motor evoked potentials) (Rothwell et al. 1987). In the presence of ongoing oscillations, the shorter the delay/latency, the higher the frequency of the oscillation. On the other hand, because CReST is a slow-tract involving multiple synapses (Jang and Lee 2019; Peterson 1979b; Peterson 1979a; Peterson, Pitts, and Fukushima 1979), the time delay of this subcortical drive is expected to be longer (something which is indeed evident in long latencies of the ipsilateral motor evoked potentials) (Taga et al. 2021; Ziemann et al. 1999); therefore, in the setting of continuous oscillatory neural coupling, the frequency should be lower than is the case for CST. Beyond these basic physiological and anatomical considerations, there is also empirical evidence that such frequency-specificity may exist. Previous work has suggested that beta (15-30 Hz) and low-gamma (30-50 Hz, i.e., Piper frequency (Brown 2000)) may originate from CST (Conway, Halliday, et al. 1995; Salenius et al. 1997; Kilner et al. 2000; Szurhaj and Derambure 2006; Brown et al. 1998; Farmer et al. 1993; Conway, Farmer, et al. 1995), at least for the fast-conducting axons (Ibanez et al. 2021). Therefore, activity in these frequency bands could be used to index cortical drive (CST). In contrast, some previous work has suggested that the alpha band (5-15 Hz) may putatively originate from CReST (Grosse and Brown 2003; Chen et al. 2018); therefore, alpha could be used to index subcortical drive (CReST) (for further details see Charalambous and Hadjipapas (2022)). Hence, characterization of oscillations in alpha, beta and low-gamma bands may provide further empirical evidence on the role of either pathway during walking. Given that collecting electromyography (EMG) during walking is relatively feasible, being able to calculate and decouple these frequencies during offline analysis, especially in individuals post-stroke, may provide a window on how these descending drives contribute to different muscle pairs functionally active during walking and consequently be targeted for restoration to promote locomotor recovery post-stroke (Jang and Lee 2017; Seo YS and SH 2020).

A few studies have so far characterized alpha, beta and low-gamma oscillations in corticomuscular or intramuscular coherence during walking, in both neurotypical (Hansen et al. 2001; Halliday et al. 2003; Petersen et al. 2012; Roeder et al. 2018; Jensen et al. 2019; Zipser-Mohammadzada et al. 2023; Kitatani et al. 2023) and neurologically impaired (Hansen et al. 2005; Norton and Gorassini 2006; Nielsen et al. 2008; Kitatani et al. 2016; Lodha et al. 2017; Weersink, de Jong, and Maurits 2022; Zipser-Mohammadzada et al. 2022) cohorts. However, most of those studies focused on a particular group of muscles, namely the ankle dorsiflexors, which are thought to be mainly CST-driven (Brouwer and Ashby 1992; Brouwer and Qiao 1995; Bawa et al. 2002). These studies showed that beta oscillations are predominantly present in tibialis anterior (TA; i.e., ankle dorsiflexor) during the swing phase and may be absent post stroke (Hansen et al. 2001; Halliday et al. 2003; Nielsen et al. 2008; Petersen et al. 2012; Kitatani et al. 2016).

While equally important as dorsiflexors in walking, plantarflexors (Soleus – SOL, Lateral Gastrocnemius – LG, Medial Gastrocnemius – MG) have received less attention in the aforementioned literature. In neurotypical adults, Clark et al. (2013) demonstrated that low-gamma band (i.e., 30-60 Hz) of SOL-MG was modulated depending on the task (i.e., reduction during dual task and increase during a unilateral long step) and was positively correlated only with the ankle plantarflexor power during the unilateral long step task; no modulation in any other bands. Also, in neurotypical adults, Jensen et al. (2019) showed that the neural coupling between the soleus (SOL) and medial gastrocnemius (MG) during stance was present in wide range (e.g., 10-20 and 25-45 Hz), yet their coherence was lower compared to that measured in dorsiflexors. In stroke patients, Kitatani et al. (2016) reported significantly lower LG-MG coherence in the beta band (15- 30 Hz) on the paretic side than in non-paretic side, but not for the low-gamma. Interestingly, they did not find any significant relationships between LG-MG coherences with any clinical measures. Furthermore, Lodha et al. (2017) demonstrated that only the gamma neural coupling of SOL-MG (i.e., 30-60 Hz) differed between healthy controls and stroke patients and this difference (i.e., control and non-paretic leg > paretic leg) depended on the walking task (i.e., long step). Taken together, both beta and low-gamma bands are present in plantarflexors during walking, low-gamma might be modulated by the task specificity, and both beta and low-gamma might be degraded after stroke.

While these few studies did also examine plantarflexors during walking, important gaps in knowledge are still present. Four important outstanding questions still exist. First, what is the involvement of the alpha band (e.g., potentially CReST) during walking? Second, what is the effect of walking speed on the common neural drive to plantarflexors? Third, what is the distribution of coherence across all three plantarflexors? Fourth, what is the functional impact of the IMCs on walking-specific neuromechanical measures that quantify muscle-specific mechanical output? Towards the first question, given that plantarflexors may receive also a subcortical drive (e.g., CReST) (Nielsen and Petersen 1995; Diete-Spiff, Carli, and Pompeiano 1967; Cowan et al. 1986), thus far, there is limited evidence on alpha band coherence during walking, in both neurotypical adults and individuals with stroke. Towards the second question, walking speed is a major factor for walking neuromechanics (Kirtley, Whittle, and Jefferson 1985; Sousa and Tavares 2012; Hof et al. 2002); so, ideally different walking speeds (e.g., slow, comfortable, fast etc.) should be analyzed. Towards the third question, given that individual plantarflexors may differ in neural and physiological properties (Tucker and Turker 2004; Kinugasa, Kawakami, and Fukunaga 2005; Cronin et al. 2013; Hug et al. 2021), there is limited IMC data across the full distribution of the three plantarflexors (i.e., SOL-LG, SOL-MG, LG-MG) during walking. Towards the fourth question, the studies that investigated the relationships between plantarflexors’ IMCs and functional measures of walking behavior used global walking measures (e.g., walking speed) (Kitatani et al. 2016; Lodha et al. 2017). However, these measures can be affected by multiple factors and are not muscle-specific. In order to better understand the linkage between the neural and mechanical control of a muscle group during walking, a muscle-specific mechanical walking measure would be more informative (e.g., ankle angular velocity can be used as a surrogate of the function of TA during the swing phase of walking) (Charalambous 2015; Srivastava et al. 2023).

In this paper we set out to explore these still outstanding research questions in a clinical cohort of chronic stroke individuals. To this end, we investigate the common neural drive (as quantified by IMC) and across a large range of frequencies (5-50 Hz) including the alpha (5-15 Hz), beta (15-30 Hz), and low-gamma (30-50 Hz) bands and across the entire distribution of plantarflexors (SOL-LG, SOL-MG, LG-MG). Also, to investigate the linkage between neural and motor control of post-stroke walking, we characterized the relationships between those IMCs and walking-specific measures namely the propulsive impulse (PI; muscle-specific mechanical output) and walking speed (behavioral). As both neural drive and biomechanical measures may change with walking speed, here we investigate IMCs and their relationships at two walking speeds: self-selected walking speed (SSWS) and fast walking speed (FWS).

## MATERIALS AND METHODS

### Ethical Approval

The present study conformed to the standards set by the latest revision of the Declaration of Helsinki. After a thorough explanation of the objectives and procedures of this study, all stroke participants read and signed a written informed consent approved by the Institutional Review Board at Medical University of South Carolina (MUSC).

### Subjects

Individuals with chronic stroke were recruited from the MUSC Stroke Recovery Research Center’s subject database. The data used and presented here were collected (but neither analyzed nor reported) as part of a larger study that examined the relationships between corticomotor responses and task-specific neuromechanics of the paretic ankle plantarflexors (SOL) and dorsiflexors (TA) in individuals post-stroke (Charalambous 2015). Inclusion criteria were: 18 to 85 years of age; ≥6 months post-stroke; either a cortical or subcortical lesion and either an ischemic or hemorrhagic type of stroke; residual paresis in the lower extremity (FM-LE motor score <34) preservation of minimal DF and PF contraction (at least 2 of 5 on a manual muscle test); ability to walk 1 minute on a treadmill with minimum speed of 0.2 m/sec; passive ROM of 5° of plantarflexion; provision of informed consent; and no cognitive impairments that made it difficult to follow the instructions about experimental procedures (3-step motor command). Exclusion criteria were: a history of seizures or used medications that could lower seizure thresholds; history of brain injury or preexisting neurological disorder; presence of pacemaker or intracranial metallic implants (Rossi et al. 2009); or severe arthritis or orthopedic problems that limited passive ROM.

#### Experimental Procedures

In this cross-sectional observational study, walking assessment was conducted in a single session, which followed a 30 min rest after a TMS session, and lasted approximately an hour. Prior to any walking trials, surface electromyography (sEMG) sensors (Motion Lab Systems; Baton Rouge, LN, USA) with inter-electrode distance of 17 mm were placed on the 3 ankle plantarflexors bilaterally using standardized guidelines (Cram and Criswell 2011). Specifically, we placed the sEMG sensors over the SOL (i.e., 2/3 of the line between the lateral femoral condyle to the lateral malleolus), LG (the most prominent aspect of the muscle belly 2 cm laterally from the midline), and MG (the most prominent aspect of the muscle belly 2 cm medially from the midline) while participants were in upright position (see the schematic in Fig 1C-I). In addition to sEMG, a reusable stainless-steel ground reference passive electrode with a diameter of 30 mm (Natus Medical Incorporated; San Carlos, CA, USA) was placed on the patella.

**Figure 1.**
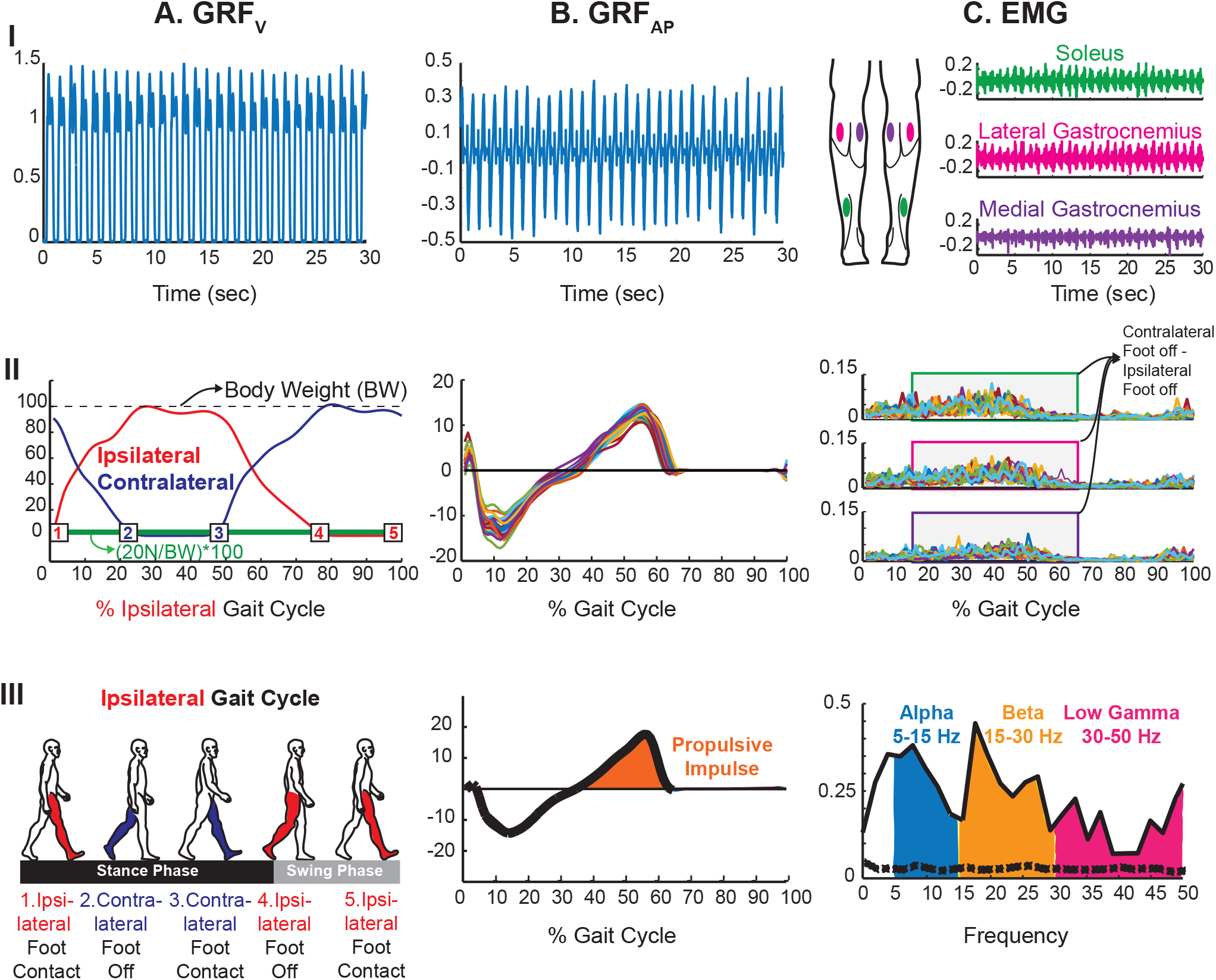
Schematic of the analysis of GRF_V_ (A), GRF_AP_ (B), and EMG (C) data. *A,* After digital processing, the raw data of GRF_V_ (I; y-axis: Volts) were used to determine the bilateral gait events (II; y-axis: % body weight). Using two ipsilateral subsequent foot contacts, a gait cycle was determined (III). The analyses of both EMG and GRF_AP_ were focused on during the stance phase (EMG: between contralateral foot off and ipsilateral foot off; GRF_AP_: second half of stance). The numbers in II correspond to the gait events in III. The green line in II denotes the threshold used to identify the gait events bilaterally. *B,* The raw data of GRF_AP_ (I; y-axis: Volts) were digitally processed, divided into strides (II; y-axis: % body weight), and used to calculate propulsive impulse (% BW.sec) during the second half of stance (III; y-axis: % body weight). *C,* Surface EMG (y-axis: mV) was collected bilateral from the three plantarflexors – SOL, LG, MG (I). After a digital processing, EMG was divided into strides from which the EMG between contralateral foot off and ipsilateral foot off (i.e., single leg stance and double limb support) was used in subsequent analyses (II). After determining coherence, the area under the curve was calculated for the alpha, beta, and low-gamma bands (III; y-axis: coherence; same analysis was done for all IMC measures). The black dashed line indicates the 95^th^ percentile of the distribution of randomly obtained spectra (i.e., level of random coherence). A*-C,* Data analysis was conducted for each 30 seconds trial of raw data and was the exact the same for both walking speed conditions (3×30sec trials per condition).

Participants walked on a treadmill at their SSWS and FWS. For each speed, participants walked three 1-minute trials, yet only the last 30 seconds of each trial were collected and subsequently analyzed. To determine SSWS and FWS, participants walked comfortably first on the treadmill without any assistance and/or assistive or orthotic device, and then each speed was determined by setting the treadmill speed initially at 0.2 m/sec. Then, we gradually increased treadmill speed by 0.05 m/sec increments until participants reported that the current speed was his/her SSWS and FWS. During the SSWS and FWS trials, at least 1-minute rest was provided to the participants; when needed, longer rests were provided (no single rest lasted more than 2 minutes). Throughout walking assessment on the treadmill (i.e., SWSS always preceded FWS), participants wore a safety harness (Robertson Mountaineering; Henderson, NY, USA) secured across the shoulders and chest. The harness was mounted to the ceiling to provide protection in the event of balance loss, and did not off-loaded any body weight. Throughout all walking trials, a licensed physical therapist closely supervised the participants to ensure safety. No form of manual support was provided during the actual data collection.

#### Data Acquisition

Data from a total of 6 trials were collected; 3 trials per speed condition. The sEMG of SOL, LG, and MG and the ground reaction forces (GRF) were collected bilaterally across all walking trials while both signals were sampled at 2000 Hz. EMG signals were filtered and amplified using an anti-alias filter of 1000 Hz and gain of 2000, respectively (Motion Lab Systems MA-300 system; Baton Rouge, LN, USA). All three components of the GRF were collected from the two independent six-degree of freedom force platforms embedded in each belt within the instrumented dual belt treadmill (Bertec Corporation, Columbus, OH, USA). All data were acquired using custom scripts in LabVIEW (National Instruments; Austin, TX, USA) and were saved automatically after each trial as coded data files on a password-protected computer for offline analysis.

#### Data Analyses

The sEMG collected from SOL, LG and MG and the anterior-posterior and vertical components GRFs (i.e., GRF_AP_ and GRF_V_, respectively) were analyzed bilaterally. All analyses were the same across trials and in each speed condition, and they were performed using custom scripts in MATLAB software (MathWorks, Inc., Natick, MA). Fig. 1 depicts the offline data analysis of GRF_V_ (A), GRF_AP_ (B), and sEMG (C).

Prior to any analysis, gait events were first detected using the GRF_V_ (Fig. 1A-I), so the signal of interest could be divided into strides. These specific gait events (i.e., bilateral foot contacts and foot offs) were determined using an automated method (Zeni, Richards, and Higginson 2008). The GRF_V_ was filtered with a low-pass 20 Hz cut-off frequency using a fourth order, zero phase lag Butterworth filter (Gottschall and Kram 2003). After digital filtering, any potential offsets were corrected, and body weight (N) was calculated using bilateral GRF_V,_ which were then normalized by the body weight and multiplied by 100. A threshold of 20 N divided by the body weight (N) multiplied by 100 was used to determine the threshold crossings in GRF_V_ trace (Fig. 1A-II). Each crossing at this preset threshold denoted bilateral foot contact (i.e., heel strike) and foot off (i.e., toe off); therefore, gait cycle (i.e., starting at the ipsilateral foot contact and ending with the immediate next foot contact of the same leg (Kirtley 2006)) was determined (Fig. 1A-III). All gait events were visually checked for accuracy (i.e., each leg was stepped on the ipsilateral belt) and removing crossovers (i.e., a stride in which leg crossed over the contralateral belt).

Then, both GRF_AP_ (Fig. 1B-I) and sEMG (Fig. 1C-I) raw signals were digitally processed. GRF_AP_ was digitally processed similarly as GRF_V_. sEMG signal of each muscle was divided by the gain (i.e., 2000) and then multiplied by 1000 to convert the units from Volts to mV. Then the signal was band-pass filtered (1-500 Hz) and band-stop filtered (156-158 Hz – to remove a 157 Hz power line peak which was detected using spectral analysis) using a third order zero phase lag Butterworth filter. The signals was then rectified (to boost the timing information of the motor unit potentials in the EMG signal) (Halliday and Farmer 2010), and finally detrended to remove any offsets. Lastly, both signals were divided into strides (GRF_AP_: Fig. 1B-II; EMG: Fig. 1C-II). Given the primary function of the plantarflexors is during the stance phase of gait cycle (Sutherland, Cooper, and Daniel 1980; Hof et al. 1992; Meinders, Gitter, and Czerniecki 1998; Neptune, Kautz, and Zajac 2001; Ounpuu and Winter 1989), the analysis of coherence and PI was limited to during stance. Specifically, the PI was calculated during the second half of the stance (Fig. 1C-III), while the coherences were quantified during the period between contralateral foot off and ipsilateral foot off (the two periods between events 2 and 4 in Fig. 1A-III); a period that denotes the single leg stance and double limb support (Kirtley 2006).

#### Muscle-specific mechanical measure

The limited work in stroke investigated the relationships between IMC and global walking measures (e.g., walking speed, step length) (Kitatani et al. 2016; Lodha et al. 2017). These measures characterize globally the walking capacity and are not either muscle-or joint-specific. Therefore, in order to understand the linkage between the common neural drive to the plantarflexors and the neuromechanics of walking, we calculated a muscle- and joint-specific measure, namely PI, which is the positive time integral of the GRF_AP_ and is the force that accelerates the center of mass (COM) forward (Neptune, Kautz, and Zajac 2001; Neptune, Zajac, and Kautz 2004). The activity of plantarflexors during the second half of stance is the main active contributor to the generation of PI (Ellis, Sumner, and Kram 2014; Neptune, Kautz, and Zajac 2001; Gottschall and Kram 2003; Liu et al. 2006). Therefore, any impairment to this muscle group manifests substantial reduction in PI generation (Turns, Neptune, and Kautz 2007). Indeed, after stroke, generation of PI on the paretic side is altered; usually paretic PI is lower than the non-paretic (Bowden et al. 2006; Turns, Neptune, and Kautz 2007; Peterson et al. 2010). To calculate PI (% BW.sec), the area under the positive curve of GRF_AP_ was calculated for each stride and averaged across strides and trials (Fig. 1B-III).

#### Coherence Analysis

Depending on the gait cycle phase of interest, previous work segmented EMG signals using predetermined fixed duration (Kitatani et al. 2016; Jensen et al. 2019) and number (e.g., 100) (Kitatani et al. 2022; Nielsen et al. 2008) of epochs. Such an approach ensures that the length and number of all epochs remains the same across trials and subjects, which is convenient for subsequent computational analyses. Yet, it does remove significant portions of data given that the duration of strides and the number of strides varies between limbs and participants, especially in clinical cohorts who have asymmetric gait. Using a fixed cycle across all participants also entails the danger of ‘contamination’ from EMG data recorded at slightly different gait phases, which makes the estimation less gait-phase specific. To avoid these limitations and because the data was retrospective in nature, we employed a slightly different approach. Within each participant, we calculated the duration of the phase during gait in which plantarflexors are typically active in neurologically healthy walking (single leg stance and double limb support) across all gait cycles (Ounpuu and Winter 1989); this calculation was performed bilaterally. Then, within each limb, we determined the epoch with the shortest duration, which was used to segment the EMG signals across all gait cycles for that specific limb. By using this approach, we limited the size of data to be removed and ensured the maximum data used within each epoch (i.e., raw data during single leg stance and early second double limb support and limb for a single gait cycle). Therefore, after data segmentation, we created a vector that included all epochs of a single muscle (e.g., paretic MG); therefore, for each participant a total of 6 vectors (∼90 sec of walking) were created for each speed condition ([SOL, LG, MG] x [paretic, non-paretic]). Given that IMC is a bivariate measure, we created three new matrices for each side: SOL-LG, SOL-MG, and LG-MG. For each signal pair, we used a Hanning window to taper the data and estimated the magnitude-squared coherence (subsequently referred to simply as ‘coherence’) at the midpoint of the time window capturing the single leg stance and double limb support (i.e., contralateral foot off to ipsilateral foot off – see Fig. 1A-III), which was determined individually as per above. The between muscles raw coherence estimates (IMC_Raw_) vary by construction from 0 to 1, whereby 0 indicates no linear relationship at the specific frequency and 1, denotes a perfect linear relationship between the two signals at that frequency (Halliday et al. 1995). To maintain the estimation of coherence at the same frequency points across participants, even though gait phases of varied somewhat in length across participants, we used a window of 1000 samples (i.e., half of the sampling rate which was also corresponded to the typical duration of the walking phase we were interested in, roughly 0.5 sec), and zero-padded the signal in case of slightly shorter signals. The resultant frequency resolution was 2 Hz. To correct the bias that may result from unequal number of segments (number of epochs) across participants we also used a z-transformed coherence measure, IMC_Z_; see equation [6] in Bokil et al. (2007). Finally, for each coherence spectrum, we also created a ‘random’ coherence spectrum. This was done by shifting the order of epochs in one of the two signals in the pair such that random coherence estimation was done based on signals that came from the same muscle recordings i.e. have roughly equal univariate properties (signal-to-noise ratio, autocorrelation and spectral density) but where the temporal pairing in the bivariate (correlation) structure is destroyed. The approach is repeated 100 times and the 95^th^ percentile of the distribution of such randomly obtained spectra serves as the level of random coherence. This approach, which is akin to epoch/segment randomization, essentially serves as a conservative estimate (see black dashed line in Fig. 1C-III) of the highest level of coherence that can be expected just randomly for a specific pair of signals, when no true relationship is present. Any difference of the true coherence estimate compared to its segment-randomized version indicates significant coherence (coherence above the random level). To this end, and to estimate significant coherences in each limb separately we derived such a measure as simply the difference between the true (IMC_Raw_) and random coherence spectrum (IMC_Delta_, the difference between the black solid and black dashed line in Fig. 1C-III). By construction this subtraction also takes care of the amount of bias that may be present given for different number of epochs/strides across participants, as the same amount of bias present both in the true (IMC_Raw_; see black solid line in Fig. 1C-III) and random estimate (see black dashed line in Fig. 1C-III).

Then, the area under the curve (AUC) for the alpha (5-15 Hz), beta (15-30 Hz), and low-gamma (30-50 Hz) bands (Jensen et al. 2018) was calculated (Fig. 1C-III) using the IMC_Raw_, IMC_Delta_ and IMC_Z_ spectra. For each participant, these resulted in estimates of coherence as (AUC as per above) for each signal pair and each frequency. It is these integral values capturing the bandlimited (as per above) coherence for each of the frequency band (i.e., alpha, beta, low-gamma) and signal pair (i.e., SOL-LG, SOL-MG, LG-MG) that they entered statistical analyses.

#### Statistical Analyses

Analyses were conducted at the group level and were the same for both walking speed conditions. Thus, for a given coherence measure, an individual participant contributed a single value for each frequency band and pair of signals. Given the small sample size, we used non-parametric statistical tests in three types of analyses (IMC measure used in each analysis): Wilcoxon rank sum test (IMC_Raw_ and IMC_Z_), one-sample Wilcoxon signed rank (IMC_Delta_), and rank correlation (IMC_Raw_ and IMC_Z_). First, to examine the effect of stroke on the common neural drive, we investigated the differences in IMC_Raw_ and IMC_Z_ between the paretic and non-paretic limb (inter-limb) using the Wilcoxon rank sum test (two-tailed). Any differences in common neural drive between the two sides would primarily indicate effect of stroke, especially on the paretic side. Second, we examined within each limb (paretic and non-paretic) which coherences were significantly above level of random coherence (i.e., chance level). This essentially looks for the significant presence (above of what is expected randomly) of any common, rhythmic neural drive to the given pair of muscles during walking in either the paretic or non-paretic limb. We used IMC_Delta_ in this test and a one-sample Wilcoxon signed rank test (one-tailed) to examine whether the median of each IMC band was significantly above 0 (whereby 0 here denotes the level of random coherence). Anything below 0 implied that there was no common neural drive to that pair of muscle. In this test, as we were interested to quantify any coherence above the random level (i.e. an IMC_Delta_ > 0), we used a one-tailed test. Third, to further understand the relationships between the common neural drive and walking measures, we examined the associations between the IMC (IMC_Raw_ and IMC_Z_) and two walking-specific measures (PI and speed) using the Spearman’s rho correlation coefficient (r_s_). Only the bands, which demonstrated significant intralimb differences (see section right above), were included. Given that these analyses were conducted for each side, the adjusted significance was set at *P* < 0.0125 (i.e., 0.05/4); correction was done within each band (two coherences and two walking measures; max 4 analyses).

We present all data as mean ± SD (1 standard deviation) and where needed we present the effect sizes of the corresponding tests. For the visualization of the interlimb and intralimb comparisons, we present the mean and median of the group and the probability density estimation using violin plots (Hoffmann 2015), which depict the probability density of the group data at different values that are usually smoothed using a kernel density estimator (i.e., smoothing method using kernels as weights). We conducted the non-parametric statistical analyses using MATLAB software (MathWorks, Inc., Natick, MA), with significance set at *P* < 0.05 unless a corrected significance level was used.

## RESULTS

Fourteen individuals with chronic stroke participated; **Table 1** presents stroke subjects’ demographics, clinical, behavioral, and walking characteristics. The majority of patients were males, were over 60 years old, were less than 3 years post-stroke, had ischemic stroke and right paresis, walked nearly 1.5 times faster in FWS than in SWSS, and walked without mobility aid. Using SSWS, Perry et al (Perry et al. 1995) post-stroke walking functional category predicted that four participants were household ambulators (< 0.4 m/sec), eight participants were limited community ambulators (0.4 to 0.8 m/sec), and two participants were community ambulators (> 0.8 m/sec). For SWSS, data from all participants were analyzed, whereas for FWS, data from 13 participants were analyzed due to a sEMG artifact in a single participant (PID 10). During all walking trials in both walking speed conditions, all participants successfully completed the experimental procedures; there were no cases of losing balance, falling, or fatiguing.

**TABLE 1.** Subject’s demographics, clinical, behavioral, and walking characteristics.

| **PID** | **GENDER**<br>**(F/M)** | **AGE**<br>**(Years)** | **TIME**<br>**POST-**<br>**STROKE**<br>**(Months)** | **STROKE TYPE**<br>**(Ischemic/**<br>**Haemorrhagic)** | **PARESIS**<br>**(L/R)** | **FMLE**<br>**(Max 34)** | **SSWS**<br>**(m/sec)** | **FWS**<br>**(m/sec)** | **FWS**<br>**(%SSWS)** | **MOBILITY**<br>**AID (Y/N)** |
| --- | --- | --- | --- | --- | --- | --- | --- | --- | --- | --- |
| **1** | F | 72 | 62 | Ischemic | Right | 27 | 0.60 | 0.90 | 150 | No |
| **2** | M | 64 | 19 | Ischemic | Right | 25 | 0.50 | 0.70 | 140 | No |
| **3** | F | 47 | 9 | Ischemic | Left | 29 | 0.90 | 1.10 | 122 | No |
| **4** | M | 47 | 10 | Ischemic | Right | 28 | 0.80 | 1.20 | 150 | No |
| **5** | M | 79 | 81 | Ischemic | Right | 28 | 0.55 | 0.70 | 127 | No |
| **6** | M | 60 | 13 | Ischemic | Right | 23 | 0.20 | 0.25 | 125 | Yes |
| **7** | M | 69 | 36 | Ischemic | Right | 26 | 0.35 | 0.40 | 114 | No |
| **8** | M | 57 | 62 | Ischemic | Right | 21 | 0.35 | 0.75 | 214 | No |
| **9** | M | 61 | 21 | Haemorrhagic | Right | 27 | 0.40 | 0.55 | 138 | Yes |
| **10** | F | 58 | 69 | Haemorrhagic | Right | 25 | 0.95 | 1.25 | 132 | No |
| **11** | M | 71 | 66 | Ischemic | Left | 30 | 0.45 | 0.70 | 155 | No |
| **12** | F | 34 | 12 | Ischemic | Right | 19 | 0.40 | 0.80 | 200 | No |
| **13** | F | 79 | 10 | Ischemic | Left | 26 | 0.35 | 0.45 | 129 | Yes |
| **14** | F | 63 | 14 | Ischemic | Right | 31 | 0.40 | 0.50 | 125 | No |
| **N = 14** | 6F | 62 ± 13 | 35 ± 27 | 12 Ischemic | 11 Right | 26 ± 3 | 0.51 ± 0.22 | 0.73 ± 0.29 | 144 ± 29 | 3 Yes |

Group data are means ± SD. PID, participant identification number; F, female; M, male; L, left; R, right; FMLE, Fugl-Meyer lower extremity, SSWS, self-selected walking speed; FWS: fast walking speed

Fig. 2 shows typical coherences of SOL-LG, SOL-MG, and LG-MG and GRF_AP_ from three participants (PID6: household ambulator; PID1: limited community ambulator; PID10: community ambulator) while they walked at their SSWS. In all three participants, the paretic coherence was generally lower than in non-paretic across all three pairs of muscles. Similarly, the paretic GRF_AP_ was generally lower than in the non-paretic. Note for PID6 (household ambulator), coherence on the paretic side was minimal across the three muscle pairs, with his GRF_AP_ mainly braking and propulsive on the paretic and non-paretic limb, respectively. In contrast, both PID1 (limited community ambulator) and PID10 (community ambulator) demonstrated some coherence across all three muscles pairs on the paretic side and more typical GRF_AP_ patterns with slightly reduced propulsion on the paretic sides.

**Figure 2.**
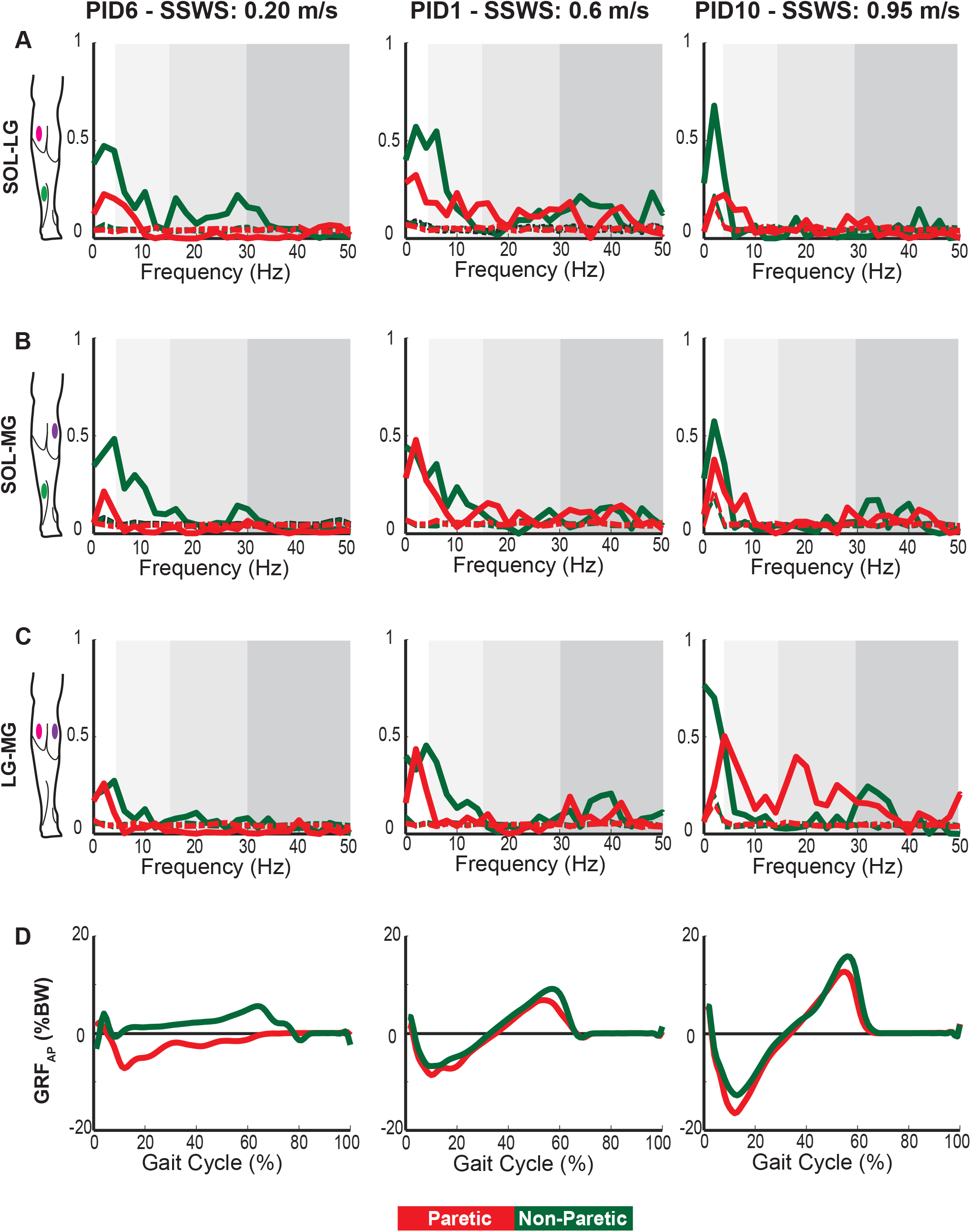
Bilateral individual IMC_Raw_ and GRF_AP_ from three subjects during SSWS. *A-C,* Solid colored lines denote the coherences at frequencies 0-50Hz calculated for SOL-LG (A), SOL-MG (B), and LG-MG (C). Colored dashed lines denote the 95^th^ percentile of the distribution of randomly obtained spectra (i.e., level of random coherence). The shaded gray boxes indicate the alpha (5-15 Hz), beta (15-30 Hz) and low-gamma (30-50 Hz) bands. *D,* GRF_AP_ is depicted bilaterally. *A-D,* Red and green lines (both solid and dashed) denote paretic and non-paretic sides, respectively.

Fig. 3 and Fig. 4 depict group (black lines) and individual subject data (colored lines) for each side. Specifically, Fig. 3 depicts the data of the paretic and non-paretic IMC_RAW_ across muscles pairs for both speeds, whereas Fig. 4 depicts the data of the paretic and non-paretic GRF_AP_ in both walking speed conditions. For both measures, there is apparent inter-subject variability, which is likely due to both the different speeds (see SSWS and FWS columns in Table 1) walked by the participants and differing levels of subject impairment, which was quantified using the Fugl-Mayer Lower Extremity (FMLE) scale (see FMLE column in Table 1).

**Figure 3.**
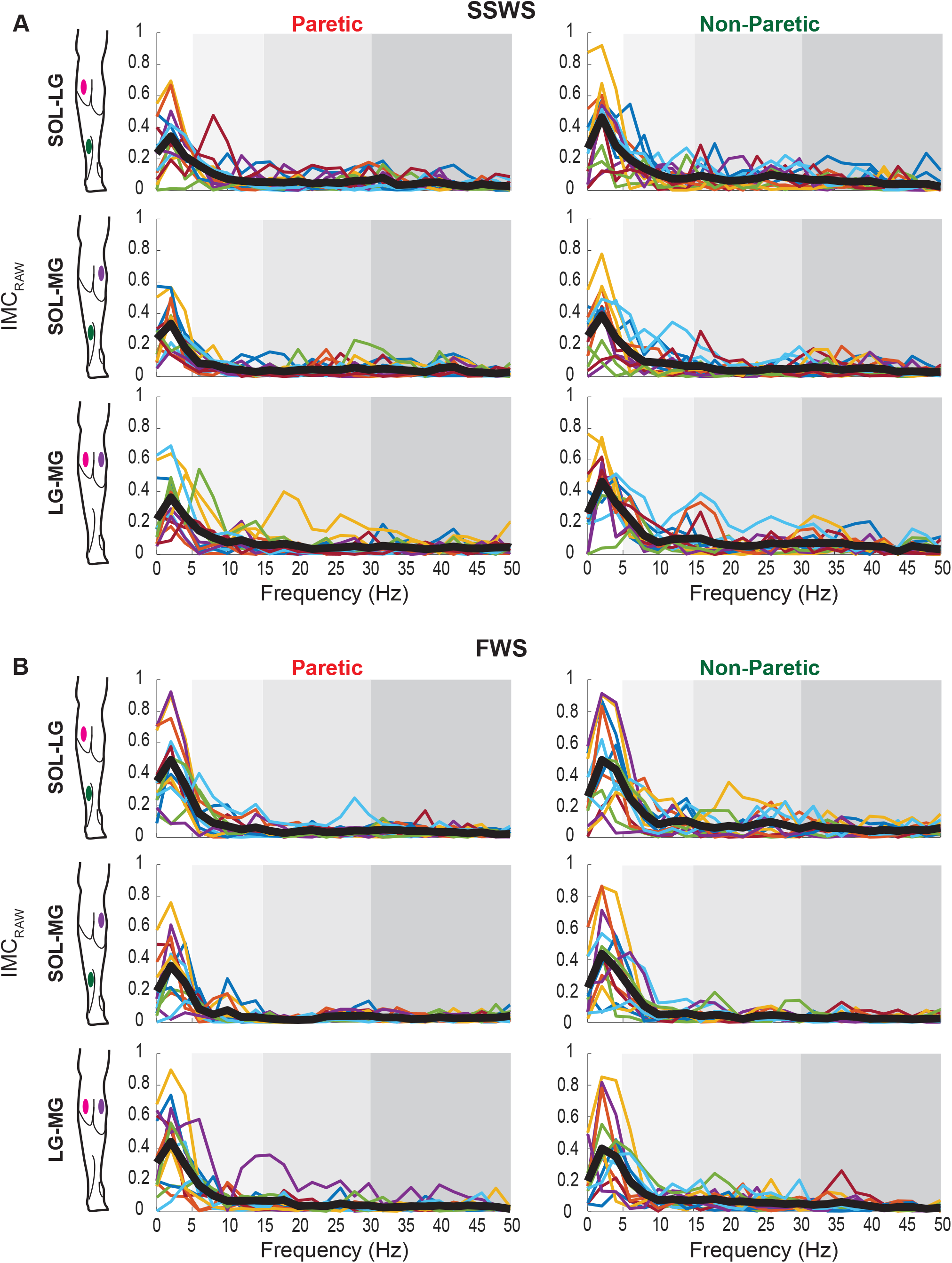
Bilateral individual and pooled coherence (IMC_Raw_) data during SSWS (A) and FWS (B). *A-B,* The colored lines denote the coherence for each subject whereas the thick black lines denotes the pooled coherence of the group. The shaded gray boxes indicate the alpha (5-15 Hz), beta (15- 30 Hz) and low-gamma (30-50 Hz) bands. Note that same color line across panels indicate the data from the same participant.

**Figure 4.**
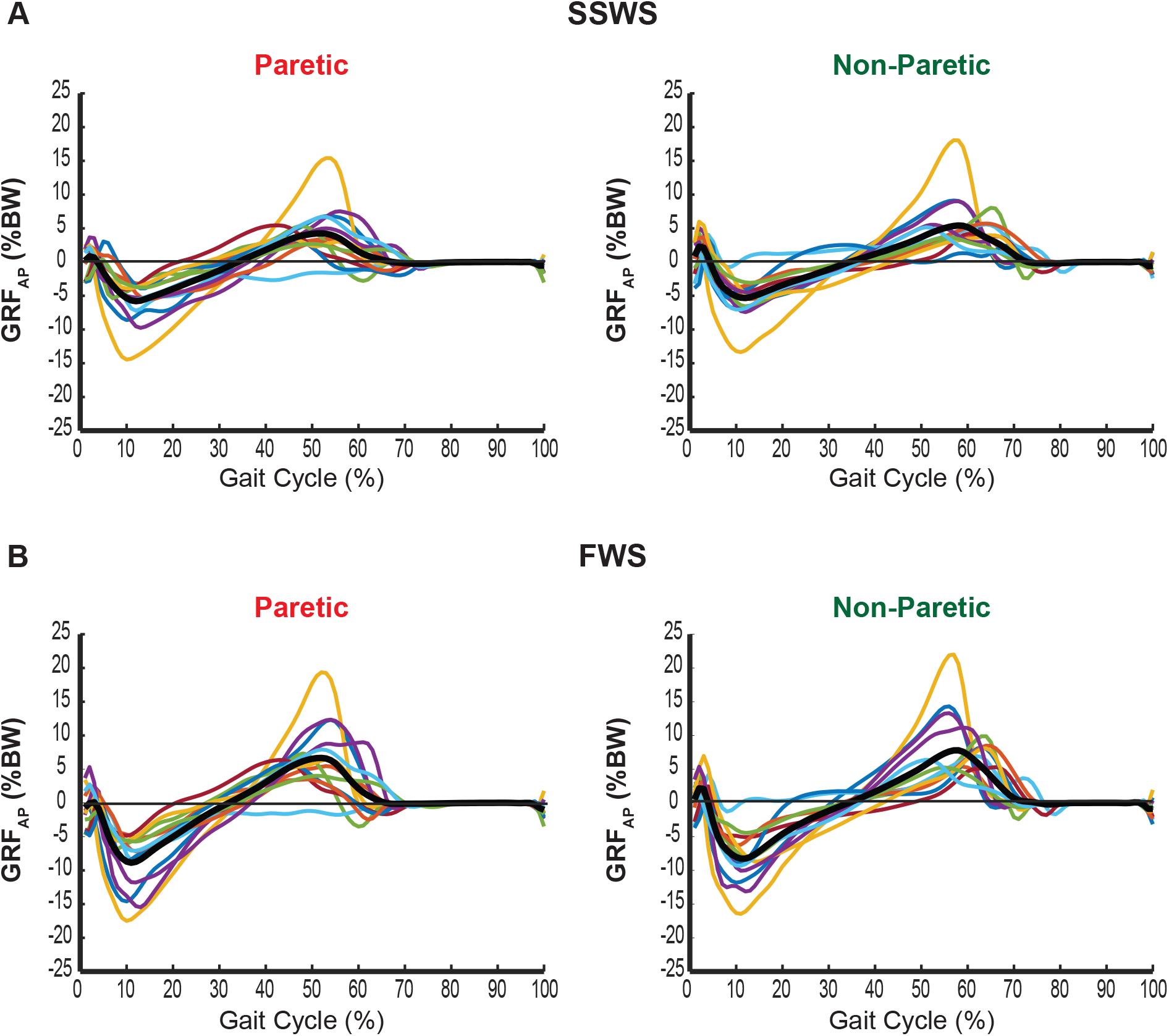
Bilateral individual and group GRF_AP_ during SSWS (A) and FWS (B). *A-B,* The colored lines denote the across trials average GRF_AP_ for each subject whereas the thick black lines denotes the average GRF_AP_ of the group. Note that same color line across panels indicate the data from the same participant.

### Differences in coherences between paretic and non-paretic side

Fig. 5 and Fig. 6 depict the interlimb differences in IMC_Raw_ and IMC_Z_ of SOL-LG (A), SOL-MG (B), and LG-MG (C) during SSWS and FWS, respectively.

**Figure 5.**
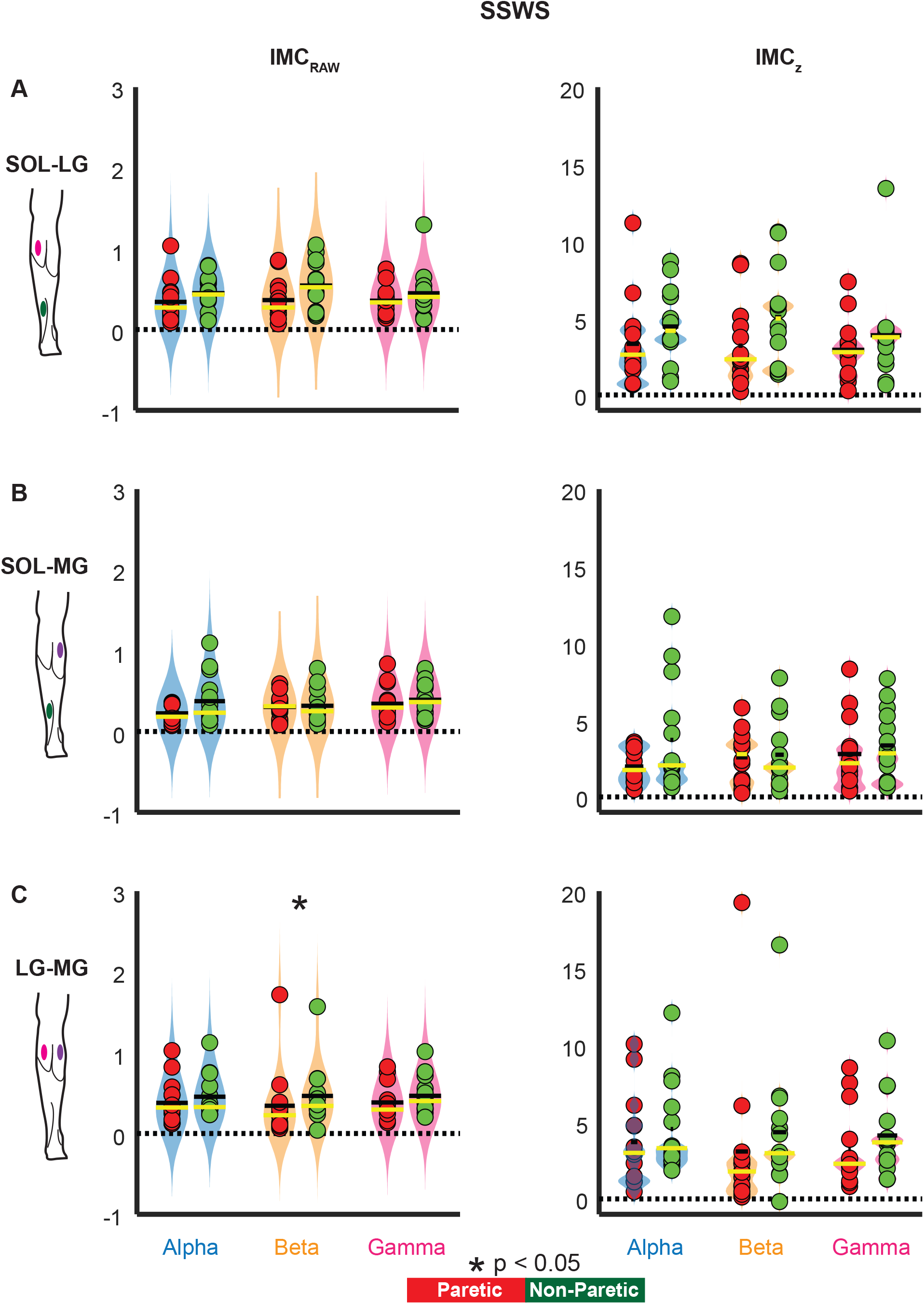
Bilateral subject and group coherence (IMC_Raw_ and IMC_Z_) of alpha, beta, and low-gamma for SOL-LG (A), SOL-MG (B), and LG-MG (C) during SSWS. Green and red dots denote individual data points (N = 14), the horizontal black and yellow lines indicate group means and medians, respectively, and the light blue (alpha band: 5-15 Hz), orange (beta band: 15-30 Hz), and light red (gamma: 30-50 Hz) shaded areas represent violin plots (see Statistical Analyses section). Black asterisk (*) denotes significance (*P* <0.05). Red and green dots denote paretic and non-paretic sides, respectively.

**Figure 6.**
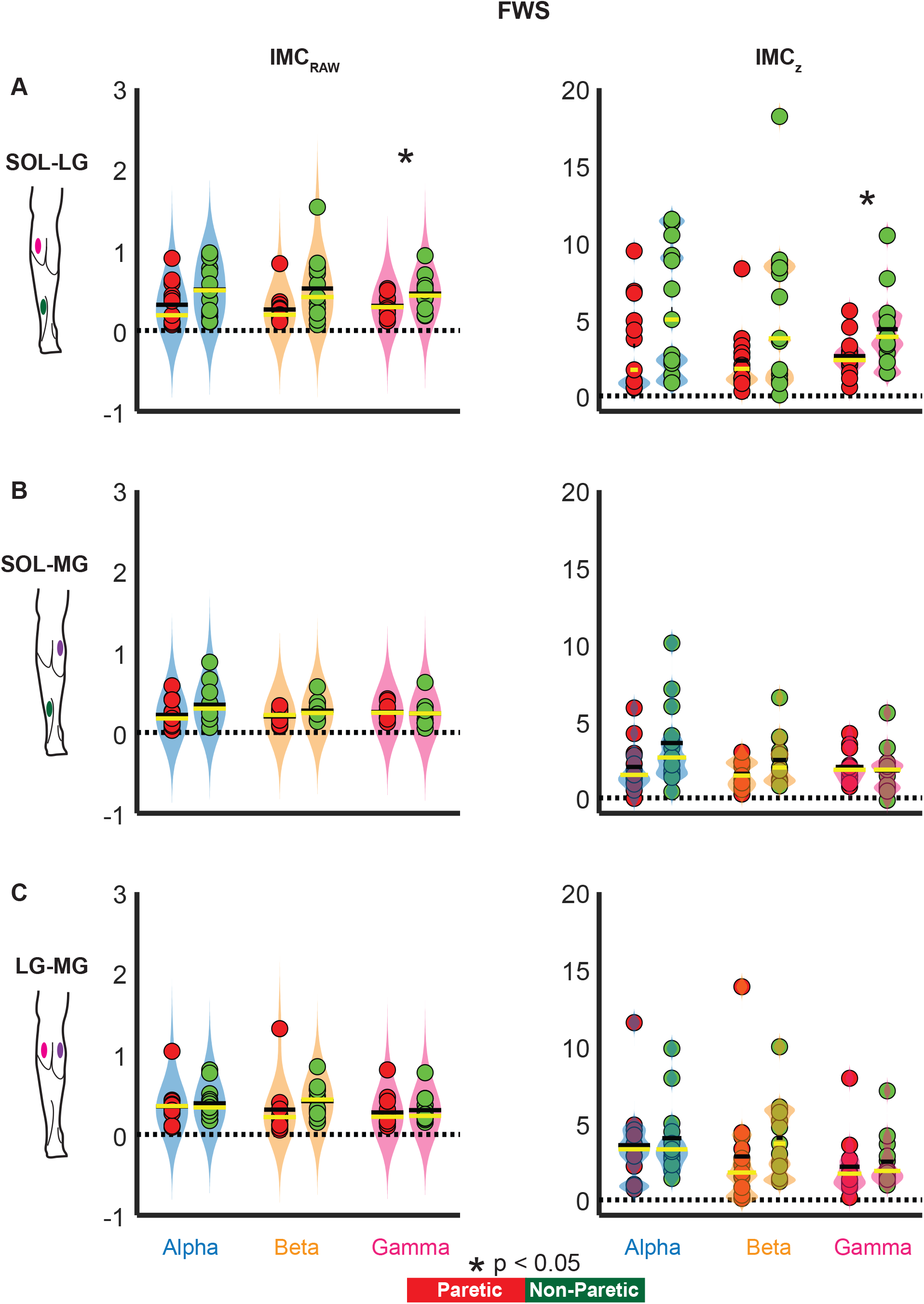
Bilateral subject and group coherence (IMC_Raw &_ IMC_Z_) of alpha, beta, and low-gamma for SOL-LG (A), SOL-MG (B), and LG-MG (C) during FWS. Green and red dots denote individual data points (N = 13), the horizontal black and yellow lines indicate group means and medians, respectively, and the light blue (alpha band: 5-15 Hz), orange (beta band: 15-30 Hz), and light red (gamma: 30-50 Hz) shaded areas represent violin plots (see Statistical Analyses section). Black asterisk (*) denotes significance (*P* <0.05). Red and green dots and bars denote paretic and non-paretic sides, respectively.

#### SSWS

Across the three frequency bands and muscle pairs, only the LG-MG beta demonstrated significant interlimb difference. Specifically, the beta IMC_Raw_ was significantly lower on the paretic side than in non-paretic side (z = -2.22; *P* = 0.0258; r = -0.42) while the beta IMC_Z_ showed a trend towards significance (z = -1.91; *P* = 0.056; r = -0.36) (Fig. 5C). For the rest of the bands and muscles pairs, no significant differences were found (all *Ps* > 0.05) (Fig. 5A-C). Note that across bands and muscles pairs, the inter-subject variability was greater for IMC_Z_ than IMC_Raw_ (i.e., longer and narrower violin plots) (Fig. 5A-C).

#### FWS

Similar to SSWS, only one band, the SOL-LG low-gamma, demonstrated significant interlimb difference. Specifically, both IMC_Raw_ (z = -2.05; *P* = 0.040; r = -0.40) and IMC_Z_ (z = - 2; *P* = 0.045; r = -0.39) of low-gamma was significantly lower on the paretic side than in non-paretic side (Fig. 6A). For the rest of the bands and muscles pairs, no significant difference was found (all *Ps* > 0.05) (Fig. 6A-C). Note that across bands and muscles pairs, the inter-subject variability was greater for IMC_Z_ than IMC_Raw_ (i.e., longer and narrower violin plots) (Fig. 6A-C).

### Differences in coherences within each side

Fig. 7 depicts the intralimb coherence IMC_Delta_ of SOL-LG (A), SOL-MG (B), and LG-MG (C) during SSWS and FWS.

**Figure 7.**
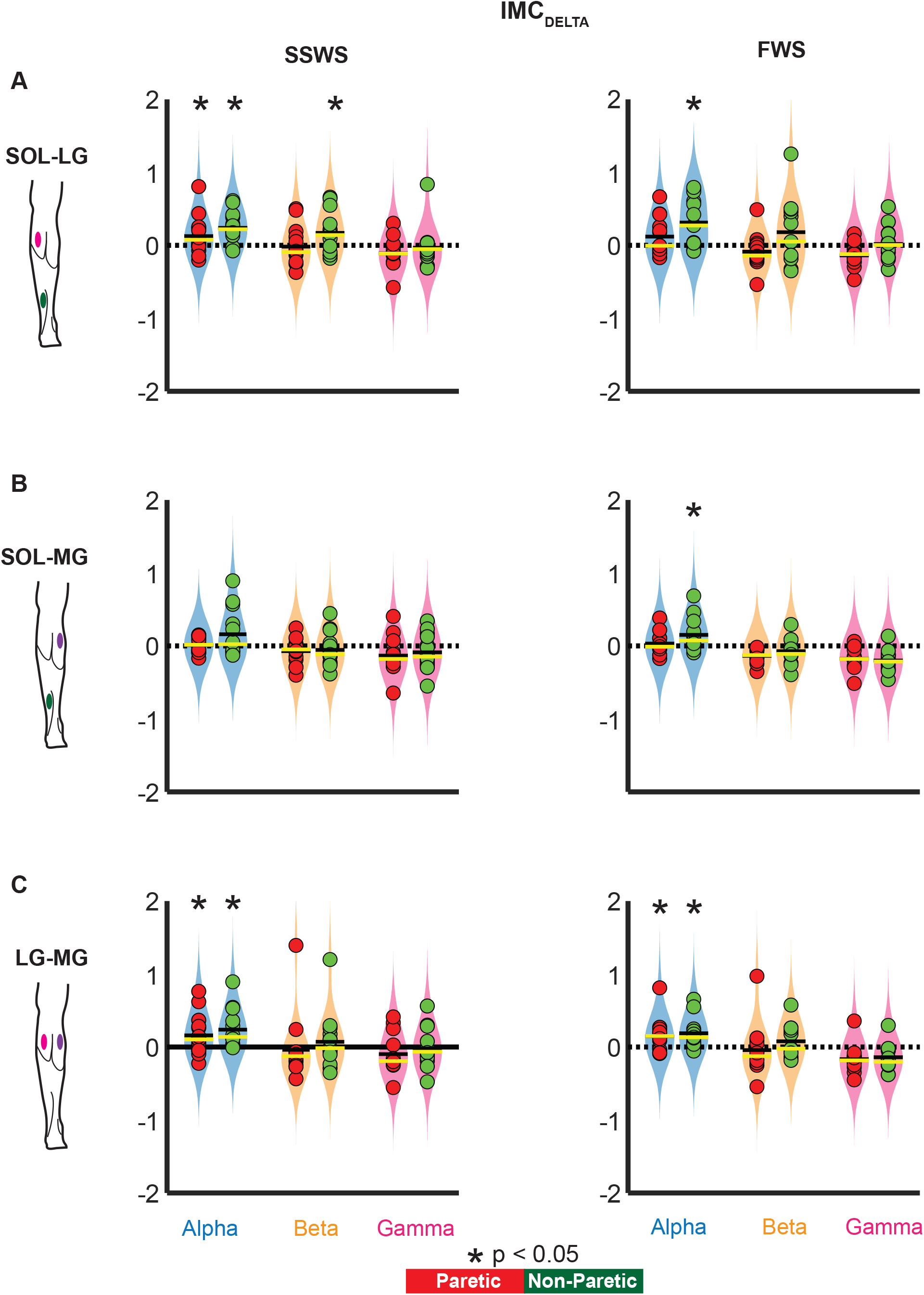
Bilateral subject and group coherence (IMC_Delta_) of alpha, beta, and low-gamma for SOL-LG (A), SOL-MG (B), and LG-MG (C) during SSWS and FWS. Green and red dots denote individual data points (SSWS: N = 14; FWS; N = 13), the horizontal black and yellow lines indicate group means and medians, respectively, and the light blue (alpha band: 5-15 Hz), orange (beta band: 15-30 Hz), and light red (gamma: 30-50 Hz) shaded areas represent violin plots (see Statistical Analyses section). Black asterisk (*) denotes significance (*P* <0.05, one-sided alternative hypothesis of Sign rank test: group median >0). Red and green dots and bars denote paretic and non-paretic sides, respectively.

#### SSWS

The group median of IMC_Delta_ of alpha was generally above 0, whereas the IMC_Delta_ of beta and low-gamma was generally below 0, with an exception of the non-paretic beta SOL-LG. On the paretic side, the IMC_Delta_ demonstrated significant intralimb differences only in alpha of SOL-LG and LG-MG. Specifically, both alpha SOL-LG (*P* = 0.045) and alpha LG-MG (*P* = 0.024) were significantly above 0. Both beta and low-gamma of IMC_Delta_ showed no significant intralimb difference in any muscle pair (all *Ps* > 0.05). On the non-paretic side, the IMC_Delta_ demonstrated significant intralimb difference only in alpha band of SOL-LG and LG-MG and beta of SOL-LG. Specifically, the alpha of both SOL-LG (*P* = 0.008) and LG-MG (*P* = 0.001) and beta of SOL-LG (*P* = 0.034) were significantly above 0. Low-gamma of IMC_Delta_ showed no significant intralimb difference in any non-paretic muscle pairs (all *Ps* > 0.05).

#### FWS

The group median of IMC_Delta_ of alpha was generally above 0, whereas the group median of IMC_Delta_ of beta and low-gamma was generally below 0, with an exception of the non-paretic beta SOL-LG (i.e., barely above 0). On the paretic side, the IMC_Delta_ demonstrated significant intralimb differences only in alpha band. Specifically, the alpha of LG-MG (*P* = 0.006) was significantly above 0. Also, on the non-paretic side, the IMC_Delta_ demonstrated significant intralimb differences only in alpha bands of all muscle pairs. Specifically, the alpha of SOL-LG (*P* = 0.016), SOL-MG (*P* = 0.011), and LG-MG (*P* = 0.001) was significantly above 0. In both sides, beta and low-gamma of IMC_Delta_ showed no significant intralimb differences in any muscle pairs (all *Ps* > 0.05).

### Correlations

Selecting just the unilateral IMCs, which were significantly above random, a total of 5 (2 paretic, 3 non-paretic) and 4 (1 paretic, 3 non-paretic) sets of correlations (4 per set) were conducted for SSWS and FWS, respectively.

#### SSWS

On the paretic side, significant strong correlations were found only between alpha LG-MG and PI. Specifically, the alpha IMC_Raw_ (*P* = 0.007; *r_s_* = 0.701) and IMC_Z_ (*P* = 0.006; *r_s_* = 0.706) of LG-MG was positively correlated with the paretic PI; greater the common neural drive was to paretic LG-MG greater the paretic PI was (Fig. 8A). No significant correlations were found between paretic alpha LG-MG and walking speed (IMC_Raw_: *P* = 0.176, *r_s_* = 0.384; IMC_Z_: *P* = 0.170; *r_s_* = 0.388) (Fig. 8B). On the non-paretic side, alpha LG-MG was not associated with either PI (Fig. 8A) or walking speed (Fig. 8B). The remaining 3 sets of correlations were not significant (all *Ps* > 0.05).

**Figure 8.**
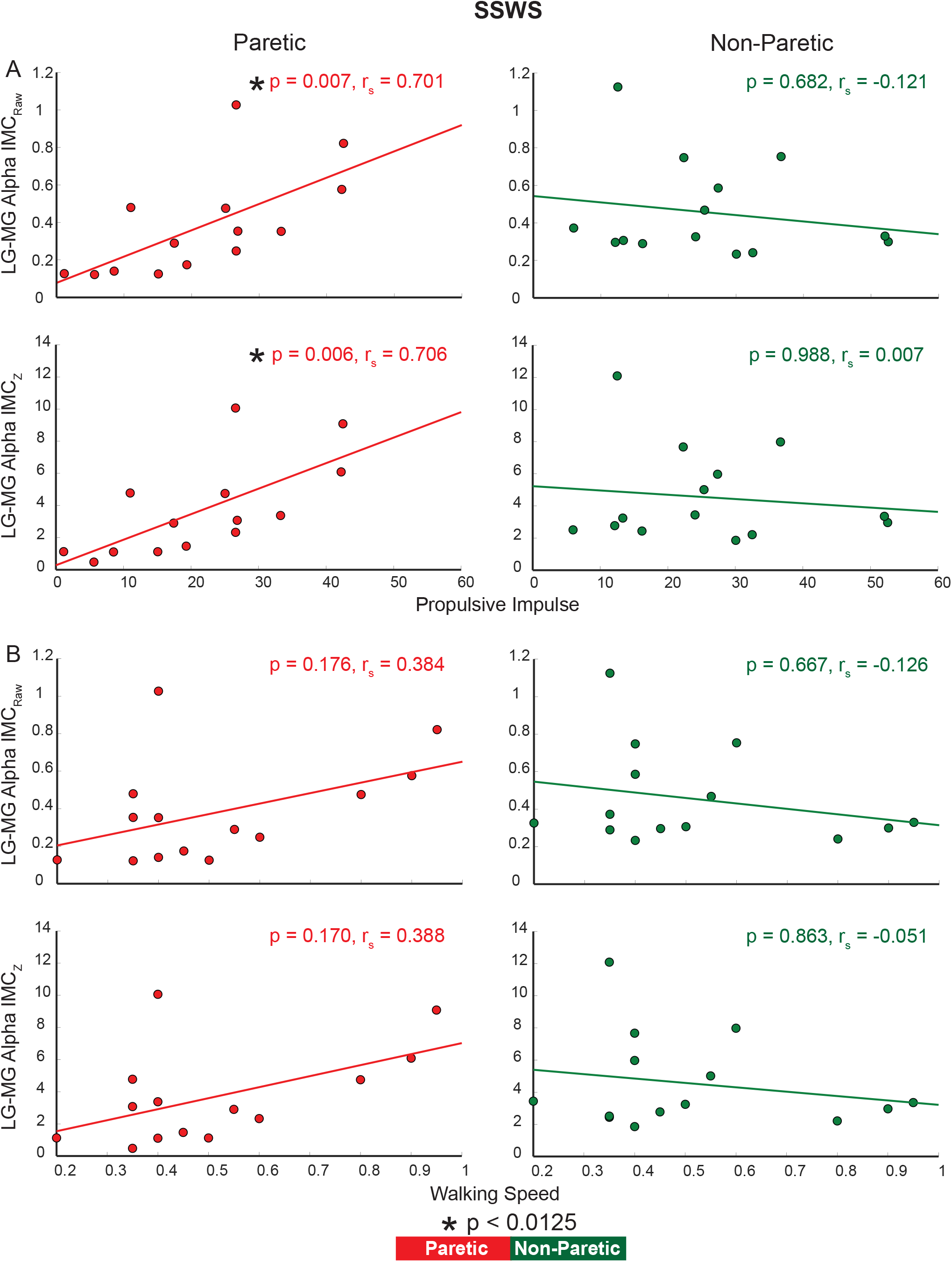
Bilateral relationships between the alpha LG-MG (IMC_Raw &_ IMC_Z_) and propulsive impulse (A) and walking speed (B) during SSWS. Green and red dots denote individual data points (N = 14) while the green and red line denote the least-squares lines. Black asterisk (*) denotes the adjusted significance level (*P*<0.0125). Red and green dots and lines denote paretic and non-paretic sides, respectively.

#### FWS

In contrast to SWSS, there was no significant correlation between alpha LG-MG and PI on the paretic side (IMC_Raw_: *P* = 0.459, *r_s_* = 0.225; IMC_Z_: *P* = 0.619, *r_s_* = 0.153). Similarly, no significant relationships were found between IMCs and walking-specific measures in any other set of correlation (all *Ps* > 0.05).

## DISCUSSION

In this study, we estimated the intermuscular coherence across the ankle plantarflexors (bilateral SOL-LG, SOL-MG, LG-MG) in the alpha, beta, and low-gamma frequency bands in chronic stroke individuals. Coherence was estimated in a time window capturing the stance phase of walking, corresponding to the phase when plantarflexors are predominantly active during walking. An instrumented treadmill was used and participants walked at two speeds: self-selected (SSWS) and fast (FWS). Our analyses revealed three main findings. First, the beta LG-MG (SSWS) and low-gamma SOL-LG (FWS) coherences were lower on the paretic side than on the non-paretic side. Second, significant alpha coherence was present bilaterally across plantarflexors (mainly non-paretic side; both speeds). Third, paretic alpha LG-MG coherence was positively correlated with paretic propulsive impulse (only SSWS). The first finding, decreased beta and gamma coherence on the paretic side only, suggests a degraded drive from the corticospinal tract post-stroke. The second finding, namely preserved alpha-coherence, suggests a possible upregulation of CReST post-stroke. The third finding, thus increased plantar flexor coherence leading to increased propulsive impulse and vice versa on the paretic side only, suggests possible contribution of the CReST to the neuromechanical function of the paretic plantarflexors during the stance phase of walking.

### Inter-limb degradation of cortical motor drive

Given that stroke may impair both CST and CReST (Paul et al. 2023), degradation of intermuscular coherence in beta and low-gamma (more related to CST) or alpha (more related to CReST) in the paretic limb was expected. Interestingly, only beta LG-MG and low-gamma SOL-LG coherences were degraded (i.e., paretic < non-paretic) in SSWS and FWS, respectively; no degradation in alpha band was present in any muscle. Taken together these findings, suggest that stroke may have predominantly impaired CST and CReST to a lesser extent, at least in our cohort.

These findings are consistent with previous work. Kitatani et al. (2016) showed significantly lower LG-MG beta coherence on the paretic side compared to the non-paretic side. In that study, however, they examined only the beta (15-30 Hz) and low-gamma (30-45 Hz) of the LG-MG pair; SOL EMG was not collected, and the alpha band was not analyzed. Also, their stroke patients walked over ground only at their self-selected speed, yet at similar speeds (0.58 ± 0.21 m/sec) as the cohort in the present study. Also, Lodha et al (2017) reported lower SOL-MG low-gamma (30-60 Hz) on the paretic side compared to the non-paretic side of post-stroke patients and to the control leg of healthy adults. Yet, this degradation was present only in the long step task and not in steady state walking in which the group preferred walking speed was 0.66 ± 0.29 m/sec. As in Kitatani et al. (2016), they examined only the SOL-MG pair; LG EMG was neither collected nor analyzed. Furthermore, they did not find any changes in the 5-13 and 13-30 Hz coherences. Despite some methodological differences across these studies and the present study, taken together they suggest that the common neural drive to plantarflexors during walking is degraded after stroke and this degradation manifests mainly in beta and low-gamma bands across speeds and walking tasks.

Such degradation has been reported in other walking muscles (i.e., TA) in individuals after stroke (Nielsen et al. 2008; Kitatani et al. 2016) and spinal cord injury (Hansen et al. 2005). These studies suggested that the degradation in those bands (e.g. beta) is due to an impaired supraspinal drive. It has been extensively reported that beta and low-gamma intermuscular coherences may be driven by CST (Conway, Halliday, et al. 1995; Salenius et al. 1997; Kilner et al. 2000; Szurhaj and Derambure 2006; Brown et al. 1998; Farmer et al. 1993; Conway, Farmer, et al. 1995); most likely from the fast-conducting axons (Ibanez et al. 2021). Beta and low-gamma band intermuscular coherence in plantarflexors during walking has been reported in neurotypical adults (Clark et al. 2013; Jensen et al. 2019). At the same time, the present study and others (Kitatani et al. 2016; Lodha et al. 2017) clearly demonstrate a degradation of these bands in plantarflexors in post-stroke walking. These findings taken together, suggest that the degradation of the neural coupling in beta and low-gamma bands across all three plantarflexors during the stance phase of walking might be due to the impaired CST originating from the contralateral/ipsilesional hemisphere. Furthermore, based on the previous work and present study, these findings strengthen the notion that the motor cortex contributes to the function of the plantarflexors during the stance phase of walking and in case of lesioned CST these contributions are impaired.

### Bilateral upregulation of subcortical drive

In order to examine the presence of any significant common, rhythmic neural drive across plantarflexors pairs during the stance phase of walking, we investigated the IMC of each pair within the paretic and non-paretic limb and compared it to a segment-randomized surrogate, which represents coherence chance detection (see methods). Across the three frequency bands, the alpha band was significantly above this random threshold in both speeds; neither beta nor low-gamma bands in any muscle pair was present in either leg (only exception was the SOL-LG beta band in SSWS). The presence of coherence in the alpha band, due to its putative relation to CReST may suggest bilateral upregulation of CReST.

Studies examining the alpha band in plantarflexors during walking either in neurotypical or neurologically impaired adults are sparse in the literature, making direct comparisons of our data with previous literature difficult. In neurotypical adults, Jensen et al. (2019) showed that SOL-MG coupling was present in wide range including 10-20 and 25-45 Hz. Although their alpha range (10-20 Hz) was slightly higher than the one used in the present study (5-15 Hz), their results showed for first time the existence of coupling in the alpha band in two plantarflexor muscles during the stance phase of walking, which is consistent with our data. We note however, that their study recruited neurotypical adults, who walked at 1 m/sec, which is higher than our clinical cohort, and that they did not collect data from LG. In our study, SOL-MG alpha coherence was significantly present only on the non-paretic side during the fast walking (0.73 ± 0.29 m/sec), while the other two plantarflexor pairs had significant bilateral presence of alpha band in both speeds.

After stroke, it has been suggested that the reliance of subcortical pathways, such as CReST, may increase due to the reduction of CST drive (Dewald et al. 1995; Turton et al. 1996). In contrast to beta and low-gamma bands, it has been previously suggested that the alpha band may putatively originate from CReST (Grosse and Brown 2003; Chen et al. 2018); therefore, alpha band intermuscular coherence could be used to index subcortical drive (CReST). It has been suggested that the major descending drive responsible for the innervation of plantarflexor motoneurons might be via oligosynaptic pathways, such as reticulospinal tract (Nielsen and Petersen 1995; Diete-Spiff, Carli, and Pompeiano 1967). Therefore, the bilateral presence of alpha band in plantarflexors in the present study may suggest subcortical contributions to that specific muscle group, which is mainly active during stance phase of walking. As the main contributors to propulsion during the second half of stance (both soleus and gastrocnemius) and to prepare for leg swing (primarily gastrocnemius), the activation of plantarflexors during late stance is crucial for executing the required mechanical tasks (Neptune, Kautz, and Zajac 2001). While CST controls the fine production of force, CReST controls for monotonic increase of force (Glover and Baker 2022). Therefore, the presence of alpha band coherence across plantarflexors in both speeds may denote a potential upregulation of CReST. Whether this upregulation is either a functional or maladaptive mechanism of recovery (Takeuchi and Izumi 2012; Jang 2013) remains to be investigated.

### Subcortical motor drive to gastrocnemius muscles is positively related to their mechanical output on the paretic side only

Most previous studies examined the association between the neural coupling, as captured by intermuscular coherence, and global walking measures (e.g., walking speed, step length, etc.) (Kitatani et al. 2016; Lodha et al. 2017). Because these measures characterize globally the walking capacity and are neither muscle-nor joint-specific, any association with muscle-specific neurophysiological measures does not adequately elucidate the linkage between the common neural drive and neuromechanics of walking. To fill this gap, we used a muscle-and joint-specific measure (i.e., PI) in those associations.

The main finding was a strong positive correlation between the alpha LG-MG and PI, yet only for this muscle group, on the paretic side, and during SSWS. Specifically, increased alpha coherence in paretic LG-MG was associated with increased paretic propulsive impulse. Therefore, the central common drive that contributes to the activation of these muscles during the stance phase of walking is related to the mechanical output of this muscle group during the same phase (Neptune, Kautz, and Zajac 2001). One interesting possibility to explain the correlation with PI existing with gastrocnemius, but not also soleus, is the role of the gastrocnemius as the primary plantarflexor contributing to propulsion of the leg into swing (Neptune, Kautz, and Zajac 2001). This differential role between the gastrocnemius and soleus muscles during walking has been demonstrated with experiments and simulations (McGowan, Kram, and Neptune 2009; McGowan, Neptune, and Kram 2008). Since leg swing initiation function is often impaired post stroke, especially in those exhibiting stiff knee gait (Brough, Kautz, and Neptune 2022), it could be advantageous to preferentially excite gastrocnemius to functionally produce both propulsion and leg swing initiation.

Interestingly, this association was present only on the paretic side; others reported strong correlation between coherence and walking kinematics only on the more impaired leg in people with spinal cord injury (Barthelemy et al. 2010). This discrepancy between the paretic and non-paretic sides in correlations may indicate that once a relatively normal level of CReST drive and PI are present, then other factors such as trailing leg angle are of increasing importance in the production of PI (Peterson et al. 2010; Liu et al. 2021; Hsiao et al. 2015; Awad et al. 2015). Neuroimaging studies have shown that disruption of CReST in the paretic leg post stroke results in poorer walking performance (Jang and Lee 2016), including reduced propulsion symmetry (Srivastava et al. 2022). Our study provides additional confirmation of the importance of CReST integrity as a functional mechanism of locomotor recovery after neurological disease (Jang et al. 2013; Jang and Kwon 2016; Jang and Lee 2017; Jang and Cho 2022).

Only few studies investigated the correlations between the neural coupling in walking muscles and *muscle-specific* mechanical output. After stroke, Palmer et al. (2021) reported that the TMS-modulated beta coherence (15-30 Hz) calculated using EEG was correlated with the paretic ankle moment; there were no correlations with FMLE (i.e., main clinical measure of motor coordination after stroke) (Fugl-Meyer et al. 1975) and walking speed. Similarly, Barthelemy et al. (2010) reported strong correlations between the intramuscular coherence of the TA and toe and knee/ankle kinematics in the more impaired limb in people with spinal cord injury. These findings taken together with ours, give rise to an important methodological insight: using a muscle-or joint-*specific* neuromechanical measure in correlations with neural coupling may feasibly unmask underlying mechanisms of the impaired motor control of walking after neurological injury. Conversely, use of *global* walking measures may mask this linkage.

The positive correlation between the alpha LG-MG and PI may also provide insight into the neural substrates responsible for the production of PI. Neurophysiological studies suggested that plantarflexors, as force producers and anti-gravity muscles, receive subcortical drive from CReST (Cowan et al. 1986; Diete-Spiff, Carli, and Pompeiano 1967; Nielsen and Petersen 1995). Given that alpha band might be a proxy of CReST, this association postulates that subcortical drive from CReST contributes to the neural origin of PI production. Yet, a few concerns on this postulation should be presented. First, association does not imply causality. Second, during locomotion, the drive from CReST commonly is mediated by spinal interneurons, which are also modulated by the central pattern generators (CPG) (Dyson, Miron, and Drew 2014). Hence, other neural substrates might be contributing to or be responsible for PI production. Third, in addition to neural inputs for the production of PI, the mechanics of the leg during walking, such as trailing limb (Peterson et al. 2010; Liu et al. 2021; Hsiao et al. 2015; Awad et al. 2015), contributes to PI. Therefore, while our data support a significant role for CReST as part of the neuromechanical process of producing PI in the paretic leg after stroke, further investigation is warranted to tease out the full extent of the contribution from CReST in relation to other neural and mechanical contributions.

The absence of any other significant correlations (i.e., muscles pairs including soleus, fast walking speed, non-paretic side), which indicate less direct contributions from CST and CReST, might be explained by the contributions of spinal networks and sensory feedback have to lower extremity muscle activity during walking (Nielsen 2003). A control system that is responsible and dominant during locomotion is the corticoreticular-reticulospinal-spinal interneuronal system (Matsuyama et al. 2004), which interacts with both CPG and sensory feedback. Furthermore, various sensory inputs (i.e., afferent feedback) might be significant drivers of plantarflexors activity during walking (Nielsen and Sinkjaer 2002). Given that the foot is in contact with the ground during the stance phase, it has been reported that positive force feedback via the Golgi Tendon Organs (i.e., force sensitive sensory organ) might be the main sensory contributor to the activity of the plantarflexors (Af Klint et al. 2009; Af Klint et al. 2010; Donelan and Pearson 2004).

### Methodological considerations

A few considerations related to methodology employed must be noted when interpreting our findings. First, we acknowledge the relatively small sample recruited for this study. To avoid Type I error, we conducted the statistical analyses using non-parametric tests and applying multiple comparisons adjustment. Second, during data acquisition, the SSWS condition always preceded the FWS condition. Though this potentially leads to fatigue, we controlled for this by keeping the walking trials short (1 min) and by providing adequate resting time between trials and also walking conditions. Third, the use of surface EMG introduces the possibility of both physiological and non-physiological sources of variability (De Luca 1997; Farina, Cescon, and Merletti 2002), particularly when quantifying the coherences during walking in both neurotypical and neurologically impaired adults (for further details see Methodological Considerations section in Charalambous and Hadjipapas (2022)). To address this concern, we adhered to the published guidelines for EMG preparation and placement (Hermens et al. 2000).

### Directions for Future Research

The main findings of this study hold significant implications for future development of a personalized rehabilitation strategy aimed at improving locomotor recovery after stroke. Given that alpha was the only frequency band positively linked with the biomechanical output of this muscle group, an intervention, either behavioral or neurophysiological, that can modulate alpha may potentially alter the walking mechanics towards more functional walking.

Such an approach is the use of transcranial alternate current stimulation (tACS), which can modulate the alpha band (De Koninck et al. 2023; Fresnoza et al. 2018). tACS is a non-invasive brain stimulation, which can target specific frequency via two potential mechanisms: either via entrainment or spike-timing dependent plasticity (Vogeti, Boetzel, and Herrmann 2022). Although tACS has already been used to boost motor performance and learning (Hu et al. 2022), only a few studies have explored the effect of tACS during walking in neurotypical adults (Kitatani et al. 2020a) and individuals with chronic stroke (Kitatani et al. 2020b). Neither of the two aforementioned studies has however, specifically targeted the alpha or beta bands, while in both studies the target muscle was TA. The main finding from both studies was that using tACS at the specific gait-frequency (single session for the neurotypical adults; 10 sessions in stroke patients) the intramuscular beta band coherence of TA during the swing phase of walking was increased. Whether this holds true with plantarflexors remains to be tested. Given that paretic propulsive impulse is impaired after stroke, entraining the motor system at alpha frequency via neuromodulation may improve propulsive impulse. Our study yields an important prediction, namely that targeting the alpha band with tACS may indeed boost both neurophysiological and muscle-specific mechanical measures in plantarflexors and subsequently promote the walking recovery after stroke.

Furthermore, numerous studies have used IMC to quantify the common neural drive to lower extremity muscles during walking in both neurotypical adults and neurologically impaired adults. Surprisingly, whether this measure is reliable and in which muscles is still under investigated. Only recently has there been effort to tackle this gap in knowledge. Zipser-Mohammadzada et al. (2023) investigated the intramuscular coherence (5-14 Hz & 15-15 Hz) of TA during two walking tasks at four various speeds in neurotypical adults; findings showed that the reliability was moderate to excellent. Future work should determine the reliability of various muscles pairs (i.e., intermuscular coherence) during walking in both neurotypical and neurologically impaired adults.

## CONCLUSIONS

In the present study, we characterized in detail, the intermuscular coherence of plantarflexors during their active phase in walking in individuals with chronic stroke. This analysis served as a valuable proxy to the descending drive to these muscles during walking. Notably, we observed a decrease in the beta and low-gamma frequency bands on the paretic compared to the nonparetic side. This suggests a degradation of the common neural drive to the paretic LG-MG (SSWS) and SOL-LG (FWS), respectively, which in turns suggests impairment of the CST. Conversely, we identified significant alpha coherence across plantarflexors mainly on the non-paretic side under both speeds, suggesting the importance of CReST drive during normal walking. Additionally, we observed significant correlations only between the alpha coherence of the gastrocnemii and their mechanical output during stance; hence, increase in propulsive impulse was associated with increased in alpha band coherence, but this was primarily evident for the paretic limb. These findings concerning the alpha coherence suggest a potential subcortical contribution to the paretic propulsion. Given that propulsive impulse is impaired after stroke, enhancing the alpha activity of plantarflexors may alleviate this impairment and facilitate walking recovery. In this vein, a frequency-specific tACS targeting the alpha frequency range could be a promising rehabilitation strategy. By targeted boosting of alpha activity in the paretic plantarflexors, this could effectively promote walking recovery by increasing propulsive impulse.

